# Dynamic methylation changes in Alzheimer’s disease-related genes during mindfulness practice – a proof-of-concept study

**DOI:** 10.1101/2025.11.17.688316

**Authors:** Idoia Blanco-Luquin, Mónica Macías, Ibai Marro, Marta Puebla-Guedea, Blanca Acha, Johana Álvarez-Jiménez, Sara Razquin-Sola, Miren Roldan, Eneko Cabezon-Arteta, Lidia González-Villena, Jesús Montero-Marín, Javier García-Campayo, Maite Mendioroz

## Abstract

**Background:** Mindfulness meditation has gained significant attention in the context of Alzheimer’s disease (AD) due to its potential effects on the prevention of cognitive decline and overall psychological well-being. The epigenetic landscape of this complex neurodegenerative disorder can be influenced by environmental factors. Mindfulness practices could actively shape the biological response to AD by modifying the underlying molecular mechanisms through epigenetic regulation.

**Methods:** A group of 17 long-term mindfulness meditators (MMs) and 17 sex- and age-matched controls were recruited. Experienced MMs participated in a 1-month Vipassana meditation retreat. Blood DNA methylation levels of differentially methylated genes (*ABCA7, ADAM10, APOE, HOXA3, NXN, TREM2* and *TREML2*) previously validated in a predictive AD-model found analyzed by bisulfite pyrosequencing.

**Results:** An increase in DNA methylation was observed for *ADAM10, APOE, HOXA3* and *TREM2* in MMs with respect to controls. Furthermore, significant differences were found for *ABCA7, ADAM10, APOE* and *HOXA3* in MMs after retreat.

**Conclusions:** DNA methylation changes identified for *ADAM10* and *HOXA3* in MMs are in the opposite direction to those occurring in AD and may constitute a protective epigenetic alteration. Mindfulness practice could counteract DNA methylation of key AD genes as a possible preventive strategy or non-pharmacological intervention in AD patients.

## Background

Mindfulness, a mental practice that consists of training the ability to focus attention on the present moment, accepting internal and external experiences without judging them [1], has been shown to have profound effects on mental and physical health [2, 3]. Its benefits include improvements in emotional regulation [4], a significant reduction in stress and anxiety [5] or the strengthening of cognitive functions such as memory and attention [6].

The relationship between mindfulness and Alzheimer’s disease (AD) is an area of growing interest, particularly with regard to how mindfulness-based therapies may mitigate cognitive decline, alleviate symptoms and influence the pathology associated with AD [7, 8]. Indeed, higher levels of trait mindfulness have been correlated with lower levels of amyloid and tau proteins, which are key biomarkers of Alzheimer’s pathology, in the brain of elders [9]. It has also been suggested that mindfulness could contribute to reduce the activity of the hypothalamic-pituitary-adrenal (HPA) axis [10], typically hyperactivated in AD. This decreased activity would favor the normalization of cortisol levels, which are elevated in the disease.

Mindfulness has been reported to reduce the adverse effects of chronic stress on the brain [11]. Research has shown that mindfulness can reduce perceived stress and improve psychological state among people with AD and their caregivers, suggesting that meditation may serve as an effective adjunctive therapy [12, 13]. In addition, the practice of meditation has been associated with improved emotional regulation and reduced anxiety, key factors in managing the psychological burden of dementia [14].

Neuroimaging studies have provided information on the structural brain changes associated with meditation. It has also been observed that mindfulness meditative practices are associated with an increase in the volume of brain regions such as the hippocampus, affected in AD, which could contribute to improved memory and attention [4, 15]. This suggests that mindfulness may enhance neuroplasticity, allowing for adaptive changes in brain structure that support emotional and cognitive health.

At the molecular level, mindfulness meditation has been linked to changes in gene expression related to chromatin modulation and inflammation. Intensive meditation retreats have resulted in rapid reductions in the expression of histone deacetylases [16]. In addition, lifestyle factors, including mindfulness practices, could profoundly impact epigenetic regulation, acting as mediators that link environmental effect to genetic expression and health outcomes. In this regard, our research group identified gene-specific methylation alterations, characterized by changes in methylation levels at distinct cytosine sites (CpG sites) across the genome, in long-term mindfulness practitioners. These changes were observed in genes associated with prevalent human diseases, including neurological and psychiatric disorders, cardiovascular diseases, and cancer [17]. This highlights the broader therapeutic implications of mindfulness as not merely a psychological intervention, but also as a potential modulator of biological pathways through epigenetic means.

On the other hand, AD has been linked to changes in epigenetic regulation, including significant alterations in methylation patterns in brain regions affected by the disease, such as prefrontal cortex [18, 19], hippocampus [20], and superior temporal gyrus [21]. AD-related DNA methylation marks are also detectable in peripheral blood and have been postulated as epigenetic biomarkers for this disease [22-24]. Our research group has developed a 6 comprehensive blood-based epigenetic panel designed to predict AD, incorporating DNA methylation markers across a diverse set of genes [25].

The purpose of this study was to evaluate whether mindfulness practice induces dynamic changes in the methylation profiles of a targeted panel of genes previously associated with AD, as assessed through blood sample analysis.

## Material and methods

### Study design and participants

A controlled, non-randomized, pre-post-intervention trial was conducted. A group of long-term mindfulness practitioners (MMs, n⍰= ⍰17) from the Spanish Association of Mindfulness and the Master of Mindfulness program at the University of Zaragoza were recruited between 3 July 2013 to 11 June 2014. Control subjects (n⍰= ⍰17) were recruited among healthy relatives and friends of the mindfulness meditators who had a similar lifestyle. Controls were matched by sex, age (± 2 years), and ethnic group, with a predominance of middle-aged men. All the participants were Caucasian (European ancestry), aged between 18 and 65 years and were fluent in Spanish. The exclusion criteria for participation were the following: having a history or current diagnosis of a psychiatric disorder based on the MINI interview, receiving pharmacological or psychological treatment, or suffering from severe medical disorders.

MMs were required to have practiced open-monitoring meditation (OM) continuously for more than 10 years before the start of the study (including the previous 10 years) with a mean of at least 60⍰min/day of formal practice during the entire period. Socio-demographic, psychological, and health-related questionnaires have been previously described elsewhere [17]. These meditators participated in a 1-month Vipassana meditation retreat at the Bujedo Monastery (province of Burgos, Spain). During the retreat, meditative practice was mainly unguided, the mean duration of daily practice was 8-9 h, the diet was vegetarian and silence was mandatory. No contact (not even by cell phone or e-mail) with the outside world was allowed [26].

### Blood samples collection and DNA isolation

EDTA plasma samples were obtained from controls and meditators before and after the retreat through venipuncture and centrifuged at 4,500 rpm for 15 min at 4°C. Buffy coat was stored at − 80°C until further use. Peripheral blood leukocytes (PBLs) DNA was isolated by the phenol-chloroform method [27]. A Nanodrop spectrophotometer (Thermo Fisher Scientific, Yokohama, Japan) was used to measure DNA concentration and purity.

### Selection of candidate methylation marks

The CpG sites selected for this study (candidate CpGs) were chosen based on previous results obtained by Acha *et al*. [25], who developed a panel of blood epigenetic biomarkers to improve prediction of AD. Candidate CpGs had been found to be differentially methylated in peripheral blood between AD patients and controls subjects and are listed in Table 1.

**Table 1.**
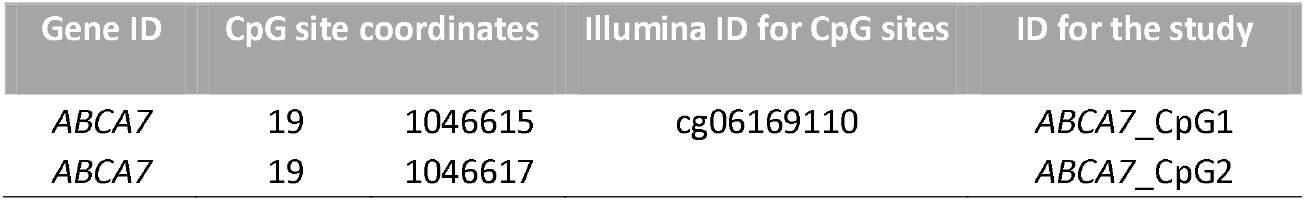
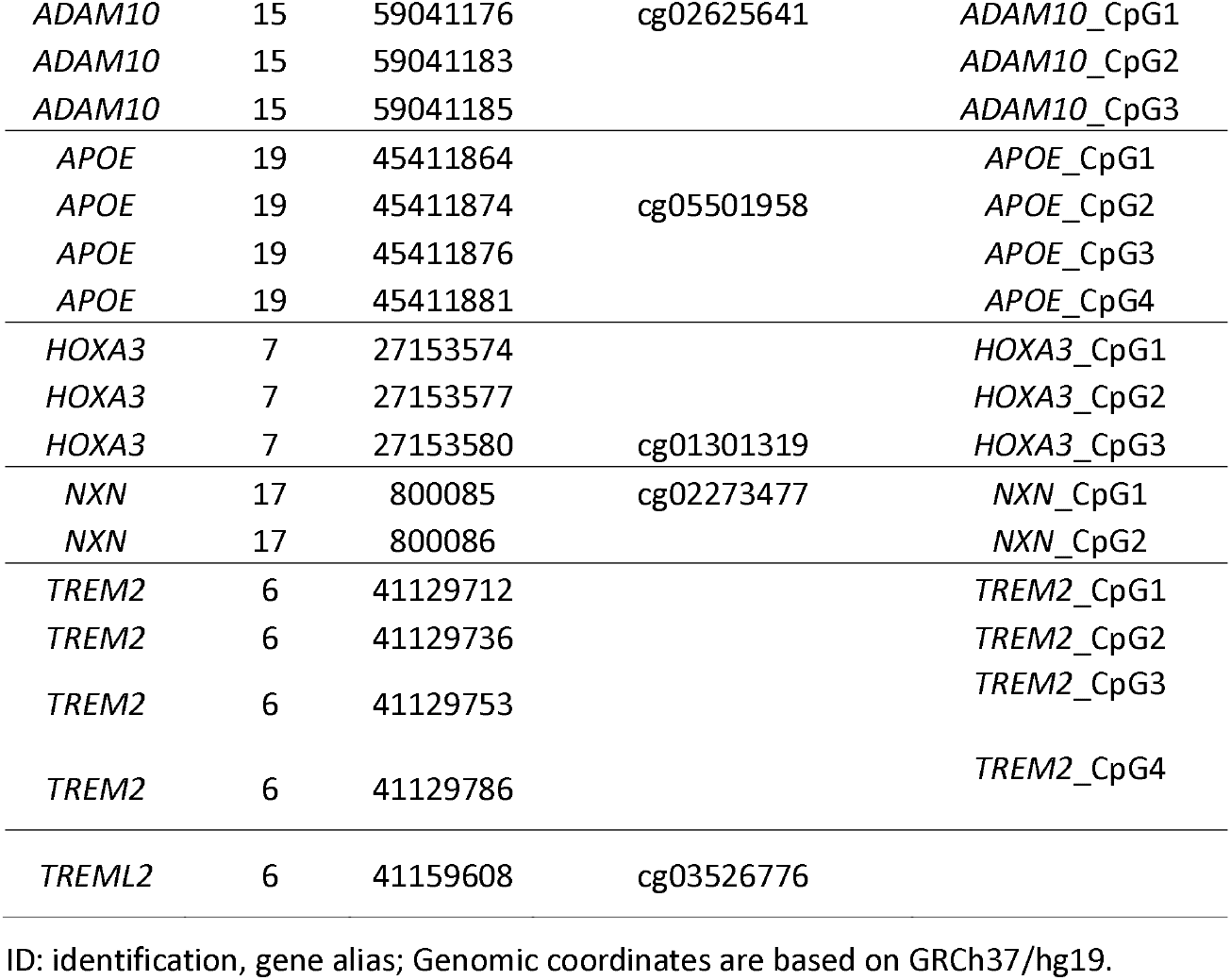
Identification and genomic position of CpG sites to be surveyed in the study.

### Determination of DNA methylation levels by pyrosequencing

Genomic DNA isolated from peripheral blood leukocytes (PBLs) (500 ng per sample) was bisulfite converted using the EpiTect Bisulfite Kit (Qiagen, Redwood City, CA, USA). Primers to amplify and sequence the target regions were designed with the PyroMark Assay Design version 2.0.1.15 (Qiagen) and were previously reported [25]. Bisulfite PCR reactions were carried out on a Veriti Thermal Cycler (Applied Biosystems, Foster City, CA). Next, 20 μL of biotinylated PCR product was immobilized using streptavidin-coated sepharose beads (GE Healthcare Life Sciences, Piscataway, NJ), and 0.4 µM sequencing primer was annealed to purified DNA strands. Pyrosequencing was performed using the PyroMark Gold Q96 reagents (Qiagen) on a PyroMark Q96 ID System (Qiagen). Unmethylated and methylated DNA samples (EpiTect PCR Control DNA Set; Qiagen) were used as controls for the pyrosequencing reaction. For each particular CpG, methylation levels were calculated with PyroMark Q96 software and expressed as the percentage of methylated cytosines over the sum of total cytosines.

### Statistical data analysis

Pyrosequencing differences in DNA methylation levels between MMs and controls and before and after retreat were evaluated according to the distribution of the continuous variables. As per Shapiro-Wilk test, for the samples that presented a normal distribution, parametric unpaired Student t-test or paired Student t-test were used, respectively, and results are expressed as mean differences (Δ) with 95% confidence intervals (CI). For samples not following a normal distribution, the non-parametric Mann-Whitney or Wilcoxon signed-rank test was used correspondingly, and results are expressed as median and interquartile ranges (IQR). Scatter plots represent the mean ± standard error of the mean (SEM) or the median accompanied by the interquartile range (IQR) for MMs vs. controls comparison. Before-after plots are used to visualize the pre-post retreat comparison in MMs. DNA methylation levels are expressed in percentages (%). Significance level was set at *p-value* < 0.05.

The SPSS 25.0 statistical program (IBM Copr., Armonk, NY) was used for the analysis of the results. GraphPad Prism software version 9.41 for Windows (GraphPad Software, La Jolla, CA, USA) was used to draw graphs.

## Results

### DNA methylation differences between long-term MMs and controls at baseline

We first examined whether long-term mindfulness meditation practice is associated with differential DNA methylation in genes previously identified as exhibiting altered methylation patterns in AD. Specifically, DNA methylation levels of 19 candidate CpGs (Table 1) across 7 genes including *ABCA7, ADAM10, APOE, HOXA3, NXN, TREM2* and *TREML2*, were investigated in peripheral blood samples of 17 long-term MMs compared with 17 controls. For this analysis, the primary variable was defined as the average methylation level of CpG sites per gene.

Average DNA methylation levels across the three CpG sites analyzed in *ADAM10* (*ADAM* metallopeptidase domain 10) were significantly higher in long-term MMs compared to controls (median: 2.960%; IQR: 2.03-4.00% vs median: 1.540%; IQR: 1.085-3.310%, *p-*value < 0.05) (Fig 1A). Each individual CpG site in *ADAM10* also exhibited increased DNA methylation levels in long-term MMs compared to controls (Fig 1B-D).

**Fig 1.**
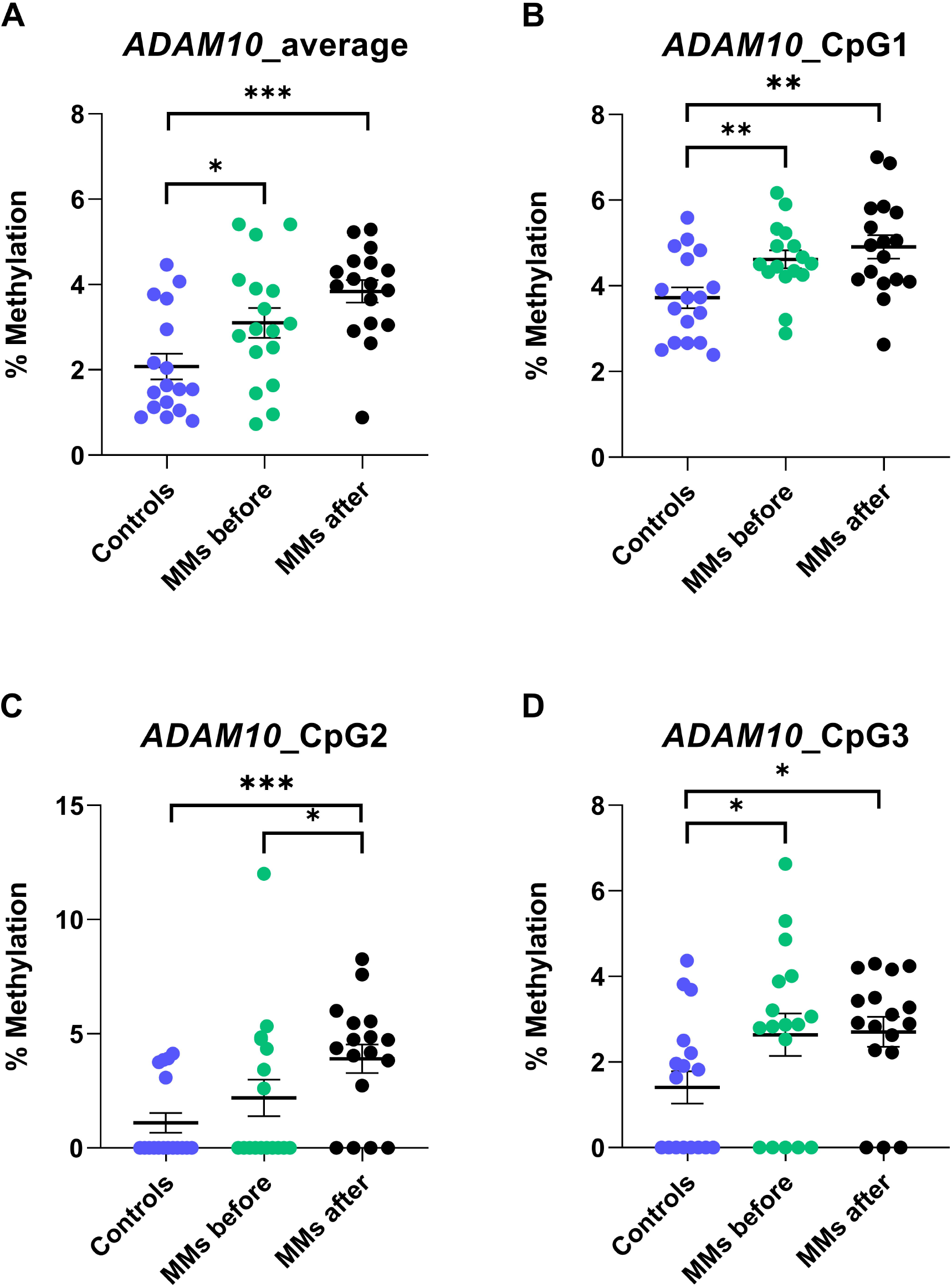
*ADAM10* methylation levels across controls and long-term MMs before and after a Vipassana-retreat. The box-plot represent the percentage of DNA methylation in average (A) and in individual CpGs (B-D) for *ADAM10* gene in PBLs measured by pyrosequencing. \**p-value* < 0.05; \*\**p-value* < 0.01; \*\*\**p-value* < 0.001.

For the *APOE* (apolipoprotein E) gene, the average DNA methylation levels resulted elevated in long-term MMs relative to controls (mean: 97.15%; CI95%: 96.58–97.72% vs mean: 96.30; CI95%: 96.02–96.58%, *p-value* < 0.01) (Fig 2). However, of the four candidate CpG sites analyzed in *APOE* only chr19: 45411881 (*APOE*_CpG4) was increased in long-term MMs versus controls (S1 Fig).

**Fig 2.**
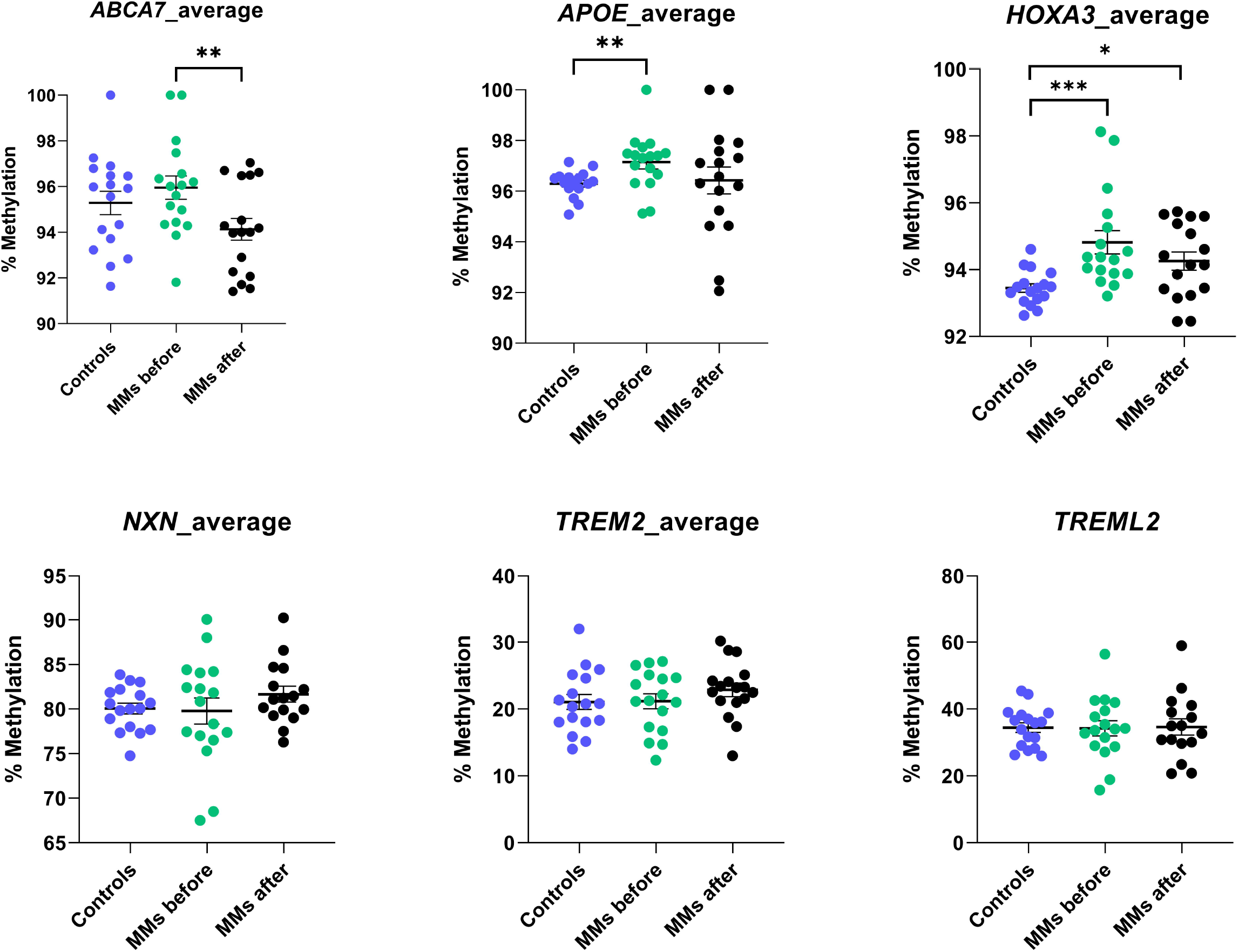
*ABCA7, APOE, HOXA3, NXN, TREM2* and *TREML2* methylation levels across controls and long-term MMs before and after a Vipassana-retreat. The box-plot represent the percentage of DNA methylation in average for all genes assayed in PBLs measured by pyrosequencing. \**p-value* < 0.05; \*\**p-value* < 0.01; \*\*\**p-value* < 0.001.

Finally, we observed a significant elevation in average DNA methylation levels for the *HOXA3* (Homeobox A3) gene in long-term MMs in comparison with controls (median: 94.38%; IQR: 93.89-95.47% vs median: 93.45%; IQR: 93.09-93.75%, *p-value* < 0.001) (Fig 2). Consistently, two of the three candidate CpG sites measured in *HOXA3* showed increased methylation levels in long-term MMs vs controls (S2 Fig).

No differences were observed for *ABCA7, NXN, TREM2*, or TREML2 when comparing long-term MMs and controls (Fig 2).

### Dynamic DNA methylation changes in long-term MMs after a 1-month Vipassana retreat

To assess the dynamic impact of intensive meditation on the selected AD-related genes, we compared DNA methylation levels in long-term MMs before and after a 1-month Vipassana retreat. This retreat is an intensive silent meditation program focused on insight meditation and mindfulness training.

We observed that average *ABCA7* (ATP-binding cassette sub-family A member 7) methylation levels significantly decreased in meditators after 1-month retreat (mean: 95.95%; CI95%: 94.87–97.03% vs mean: 94.13%; CI95%: 93.12–95.14%, *p-value* < 0.01) (Fig 2). Accordingly, *ABCA7*_CpG2 (chr19:1046617) showed a reduction in DNA methylation levels after intensive mediation practice (S3 Fig).

Following a 1-month meditation retreat, average *ADAM10* methylation levels raised compared to controls (median: 2.92%; IQR: 2.25-3.83% vs median: 1.64%; IQR: 0.00-2.36%, *p-value* < 0.05) (Fig 1A). Consistently, each individual CpG site in *ADAM10* also showed increased DNA methylation levels in meditators after retreat compared to controls. Of note, the methylation levels in *ADAM10* showed a dose-response effect across controls, pre-retreat long-term MMs, and post-retreat long-term MMs (Fig 1B-D).

In the case of the *HOXA3* gene, average DNA methylation levels remained elevated in meditators compared to controls after the retreat (median: 94.26%; IQR: 93.68-94.84% vs median: 93,45%; IQR: 93.18-93.71%, *p-value* < 0.05) (Fig 2) and in *HOXA3*_CpG3 (cg01301319) (S2 Fig). However, a significant reduction was observed among long-term MMs for *HOXA3*_CpG2 (chr7:27153577) when comparing methylation levels after and before retreat (S2 Fig).

Finally, although average DNA methylation levels in *TREM2* remained unchanged after intensive meditation (Fig 2), we observed that *TREM2*_CpG4 (chr6: 41129786) exhibit a dose-response effect similar to that observed in *ADAM10* (S4 Fig).

For *APOE, NXN* and *TREML2*, no differences in average DNA methylation levels were identified after the 1-month retreat (Fig 2).

## Discussion

In this study, we examined whether mindfulness meditation affects DNA methylation in genes previously identified as differentially methylated in AD. Using a candidate-gene approach, we observed that several AD-related genes, including *ADAM10, APOE* and *HOXA3*, exhibited distinct methylation patterns in meditators compared with non-meditators subjects. In addition, we also found a dynamic response in the DNA methylation profile, most consistently in *ADAM10*, following intensive meditation practice during a 1-month retreat. Together, these results reveal that while some AD-related genes maintain stable DNA methylation profiles, other genes, particularly *ADAM10*, exhibit dynamic DNA methylation changes in response to intensive meditation practice. This supports the hypothesis that mindfulness practice can induce both enduring and flexible epigenetic modifications in pathways relevant to neurodegeneration.

The emerging link between mindfulness and epigenetic regulation, particularly DNA methylation, may help clarify the molecular pathways through which contemplative practices modulate processes such as inflammation, oxidative stress, or immune function [28]— mechanisms that are also central to AD pathophysiology. In this regard, several studies have evaluated the association between methylation and mindfulness. Our group previously explored genome-wide DNA methylation by the Illumina HumanMethylation450 platform in the 17 long-term MMs and 17 matched controls and found 64 differentially methylated regions (DMRs), corresponding to 43 genes involved in lipid metabolism and atherosclerosis signaling pathway and linked to common human diseases, such as neurological and psychiatric disorders, cardiovascular diseases, and cancer [17]. We also demonstrated that long-term MMs showed differential DNA methylation at specific genomic *loci*, including those associated with telomere maintenance and stress response pathways [29]. In accordance, Dasanayaka *et al*. highlighted a positive correlation between meditation duration and the expression of genes related to telomere dynamics [30]. Furthermore, to evaluate the methylome of experienced meditators after a one-day intensive mindfulness practice, Chaix *et al*. performed an Illumina 450K methylation array and identified 61 differentially methylated sites (DMSs), implicating some pathways related to the immune system and DNA damage response [31]. Pragya *et al*. also found changes in DNA methylation at 450 CpG sites after an 8-week meditation program in healthy, meditation-naïve college students [32]. Pathway analysis revealed that these genes were enriched in metabolic and brain functions signaling pathways.

In the present study, we found that long-term MMs had generally higher levels of methylation in the surveyed genes compared to the controls. It is also interesting to note how the magnitude of these changes is sometimes greater when comparing controls and long-term MMs after 1-month retreat, revealing a putative dose-response effect, particularly evident for *ADAM10* gene. This observation may reflect a potential protective or compensatory epigenetic mechanism associated with meditation practice in AD-related genes.

In our study, *ADAM10* has shown the most consistent pattern of methylation changes in long-term MMs. *ADAM10* encodes the principal α-secretase responsible for non-amyloidogenic cleavage of the amyloid precursor protein (APP) [33], which prevents the formation of amyloid β (Aβ) peptides, a central pathological feature of AD. Moreover, *ADAM10* is a key regulator of neuroinflammation in the central nervous system, including the astroglia-mediated inflammatory response induced by AD pathology [34]. In mild AD, there is a displacement of ADAM10 from the membrane to the extracellular space, resulting in higher levels of the soluble, but inactive form, in plasma and cerebrospinal fluid (CSF) [35], and higher levels of soluble ADAM10 are associated with decreased cognitive performance, especially in *APOE*e4 carriers [36]. In our work, we have observed how DNA methylation levels increased in several individual CpGs located in the promoter region of *ADAM10* and in average when comparing 15 MMs and controls. Moreover, we show how this difference increases even more in long term MMs after retreat. Of note, *ADAM10*_CpG2 (chr15:59041183) had been found to be hypomethylated in AD patients in the study by Acha *et al*. [37], consistent with the elevation at protein level observed in other studies [35, 36]. Thus, *ADAM10* hypermethylation observed in long-term MMs potentially represents a protective epigenetic change in this relevant gene for AD pathology.

In this work, the *ABCA7* gene showed a decrease in DNA methylation levels in long-term MMs after the retreat. This gene encodes a transmembrane protein involved in cellular lipid transport, as well as the lipidation of apolipoproteins, including APOE. In addition, it is involved in the processing and deposition of Aβ, which directly links it to AD [38]. Consistent with our results, cg24145486 (Chr19:1054884), also located in the *ABCA7* gene, about 8,000 bp away from the CpG studied in our work (cg06169110), was shown to be hypomethylated in experienced meditators after an intensive day of mindfulness-based practice [31].

Interestingly, significant changes in *ABCA7* methylation occurred with intensive practice, but they are not observed in long-term MMs compared to controls. On the other hand, Acha *et al*. identified, at the same position, hypomethylation of the *ABCA7* gene in AD patients with respect to controls [37]. Therefore, in our hands, the sense of change does not support the idea that mindfulness meditation may have a protective effect on the epigenetic modification of this gene.

*APOE* ε4 (Apolipoprotein E epsilon 4) allele is the major genetic risk factor for sporadic AD. *APOE* gene influences AD neuropathology through multiple pathways, including altered lipid metabolism, impaired Aβ clearance, increased neuroinflammation, and synaptic dysfunction [39]. Our study showed DNA hypermethylation levels in average in long-term MMs compared 16 with controls and at chr19:45411874 (*APOE*_CpG2) in post-retreat meditators. For this cg, Acha *et al*. also reported hypermethylation in AD patients with respect to controls [37]. In this scenario, we hypothesize no neuroprotective effect of mindfulness practice on the methylation patterns with regards to this gene.

*HOXA3* belongs to the Homeobox gene family, mainly involved in early development and cell differentiation. Recent studies have concluded that homeobox genes encode transcription factors essential in neuronal function, such as Ank2-XL, an axonal microtubule organizer protein required for synaptic stability [40]. In addition, *HOXA3* gene has been implicated in several physiological processes, including inflammation and immune response [41]. Chen *et al*. also described that DNA methylation of *HOXA3* is related to the efflux capacity of high-density lipoprotein (HDL) cholesterol [42], suggesting that alterations in the expression of this gene could impact lipid metabolism. In our work, an increase of average DNA methylation levels was detected in MMs with respect to controls. Interestingly, hypomethylation was found in *HOXA3* (Chr7:27153577) in post-retreat MMs, the same position that Acha *et al*. had previously identified as hypermethylated in AD patients [37]. This decrease in methylation levels in the *HOXA3* gene could imply a beneficial effect by counteracting the methylation changes that occur in the development of AD. On the other hand, it highlights the dynamic changes in methylation that might occur for an individual meditating over the long-term compared to those that occur when the practice is intensive over shorter periods of time. This pattern could represent a positive response to a low stress environment and prolonged mental activity, contributing to a neuroprotective state.

Finally, *TREM2* gene, predominantly expressed on microglia, mediates key functions such as phagocytosis of apoptotic cells and debris, which are crucial in the context of Aβ plaque formation and tau pathology [43, 44]. Here, we found a possible dose-response effect for a CpG site located in *TREM2* (*TREM2*_CpG4, chr6:41129786), whereas the work of Acha *et al*. detected hypomethylation but in adjacent CpG sites (chr6:41129712 and Chr6:41129736). Nevertheless, we observed no significant changes in average *TREM2* methylation levels, indicating that its epigenetic regulation may be more stable and trait-like in nature.

On the whole, these findings support the notion that long-term and intensive mindfulness practice could influence genes related to AD pathogenic processes, such as *ADAM10* and *HOXA3*, through epigenetic mechanisms, specifically DNA methylation. This stable but potentially modifiable alteration could be behind certain transcriptional or structural changes observed in meditation practitioners. Further studies confirming mindfulness meditation’s ability to modify AD-associated gene methylation and expression could support the use of meditation as complementary therapies for the disease.

This study has several limitations that need to be considered. Although our sample size is limited, these findings are consistent with previous evidence suggesting that meditation and related mind–body interventions can modulate epigenetic marks involved in stress response, immune regulation, and neuronal plasticity. Future studies in larger, independent and longitudinal cohorts will be necessary to determine whether these methylation differences translate into measurable protection against neurodegenerative changes.

In conclusion, the interaction between mindfulness, epigenetics and neurodegenerative diseases underscores the importance of considering lifestyle factors along with genetic susceptibility, as interventions such as mindfulness could ultimately modulate gene expression and mitigate the risk of these pathologies. The fact that this practice has been shown to improve emotional regulation, reduce anxiety and depression, and promote cognitive health, could indicate a possible role in reducing risk factors associated with AD. Because of its remarkable impact on epigenetic regulation, mindfulness meditation could constitute a potential tool in the prevention and treatment of AD.

## List of abbreviations

AD: Alzheimer’s disease
ANOVA: analysis of variance
*APOE* ε4: ε4 allele of the APOE gene
APP: amyloid precursor protein
CpG: cytosine-phosphate-guanine dinucleotide
p-tau: hyperphosphorylated tau.

## Acknowledgements

We want to kindly thank the subjects that generously participated in the study, the Department of Psychiatry at the University of Oxford, UK, and the Spanish CIBER of Epidemiology and Public Health (CIBERESP CB22/02/00052; ISCIII) for their support.

## Funding

The project has received funding from the Network for Prevention and Health Promotion in Primary Care (RD12/0005/0006) grant from the Instituto de Salud Carlos III of the Ministry of Economy and Competitiveness (Spain), co-financed with European Union ERDF funds. The funding source did not have any influence on the design of the study, data collection and analysis, or writing of the manuscript. M.M. (Mónica Macías) is beneficiary of a Navarrabiomed postdoctoral research grant (2023-2026). J.Á.-J. has received a Doctorandos industriales grant for 2023–2026 founded by the Department of Industry and Health of the Government of Navarra. J.M.-M. has a ‘Miguel Servet’ research contract from the ISCIII (CP21/00080).

## Authors contributions

Conceptualization: Idoia Blanco-Luquin, Mónica Macías, Javier García-Campayo, Maite Mendioroz

Formal analysis: Ibai Marro, Johana Álvarez-Jiménez, Sara Razquin-Sola

Investigation: Blanca Acha, Miren Roldan, Eneko Cabezon-Arteta, Lidia González-Villena

Resources: Marta Puebla-Guedea, Javier García-Campayo

Supervision: Jesús Montero-Marín, Javier García-Campayo, Maite Mendioroz

Writing – original draft: Idoia Blanco-Luquin, Mónica Macías

Writing – review & editing: Marta Puebla-Guedea, Jesús Montero-Marín, Javier García-

Campayo, Maite Mendioroz.

Project administration: Maite Mendioroz

Funding acquisition: Javier García-Campayo, Maite Mendioroz

## Ethics approval and consent to participate

All procedures performed in this study (number PI13/0056) involving human participants were in accordance with the ethical standards of the Aragon Ethics Regional Committee and with the 1964 Helsinki Declaration and its later amendments or comparable ethical standards. All participants provided their written informed consent before participating in the study.

## Competing interests

J.M.-M. is associated with the University of Oxford Mindfulness Research Centre.

## Notes

### Competing Interest Statement

I have read the journal's policy and the authors of this manuscript have the following competing interests: J.M.-M. is associated with the University of Oxford Mindfulness Research Centre.

