## Supplementary figures and images for "Dynamic methylation changes in Alzheimer’s disease-related genes during mindfulness practice – a proof-of-concept study"

### S1 Fig

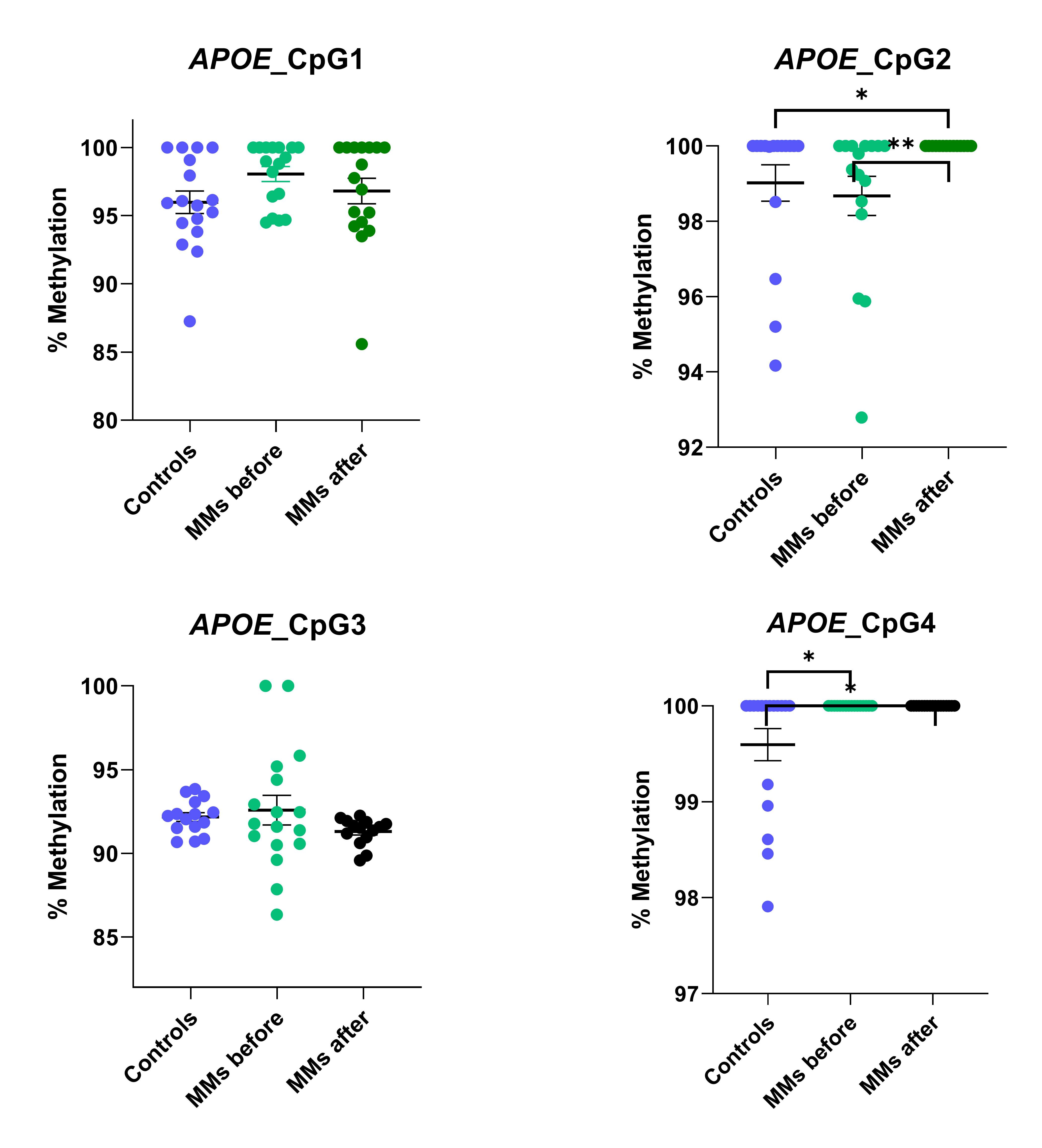

### S2 Fig

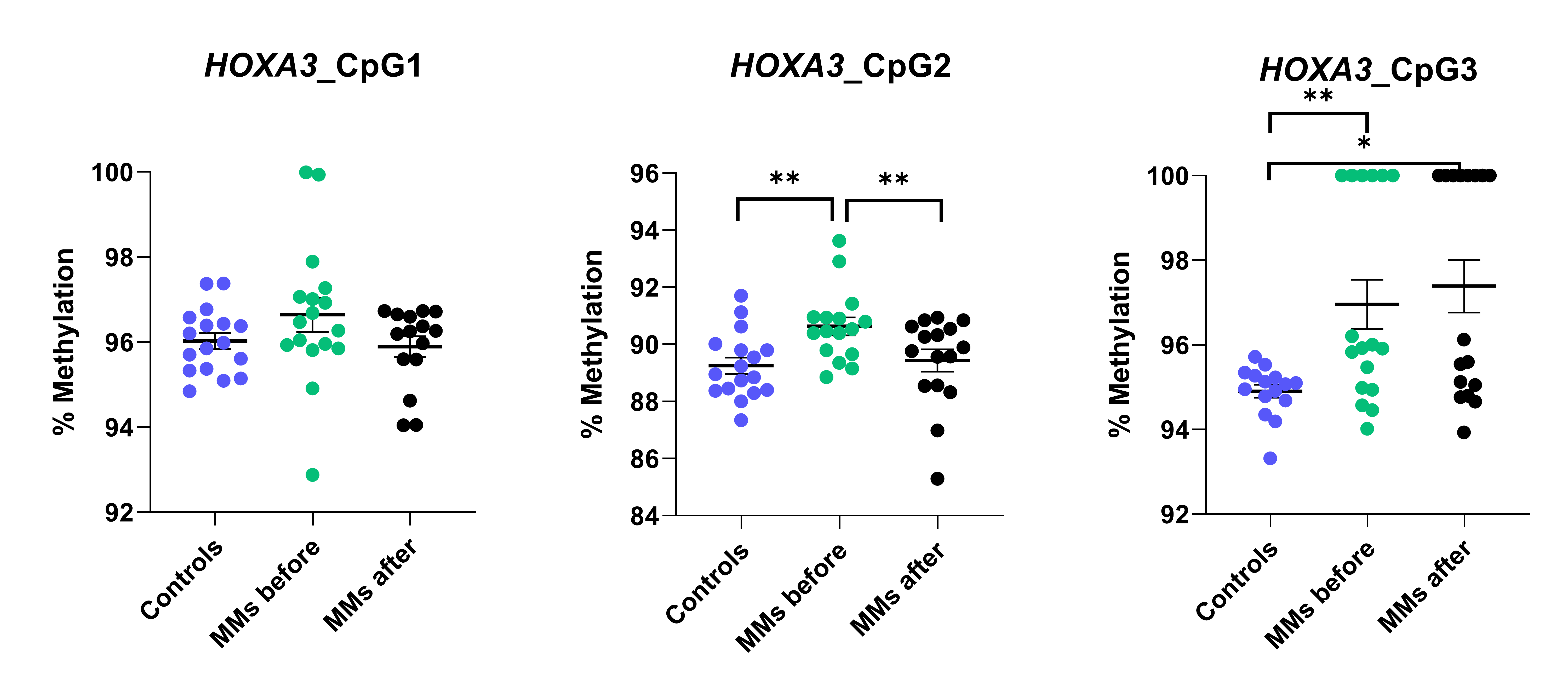

### S3 Fig

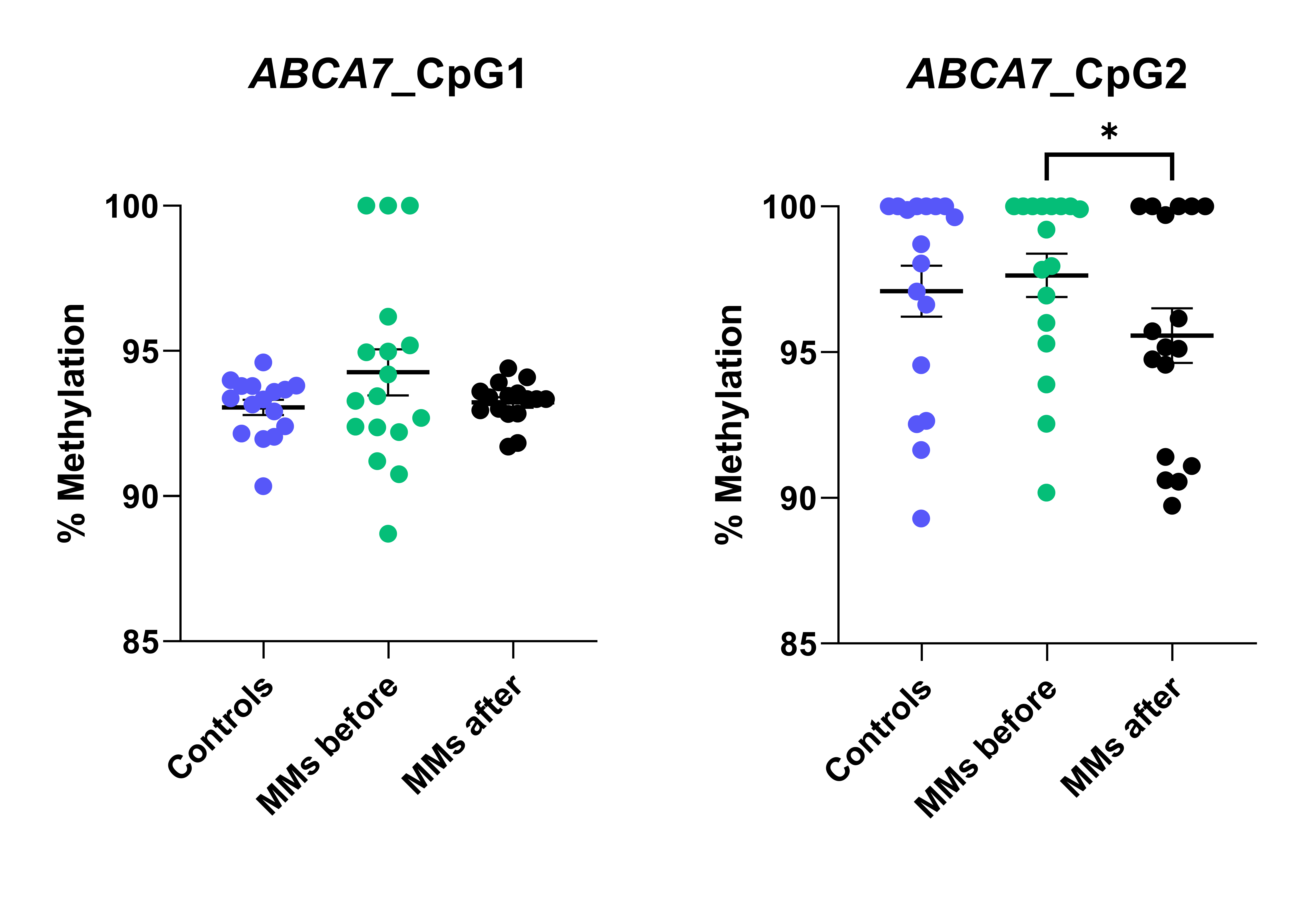

### S4 Fig

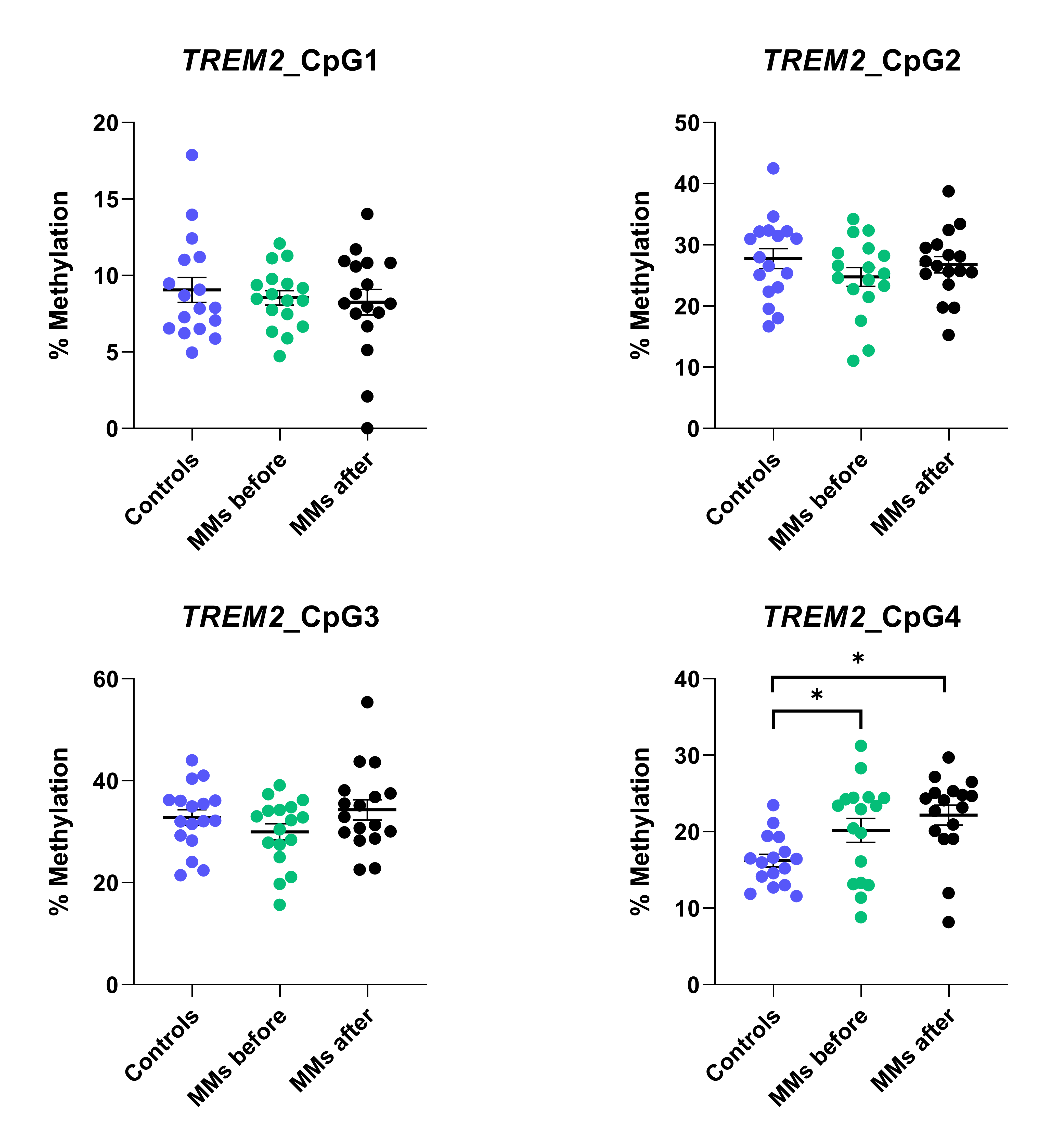
